# Comparing acute IOP-induced lamina cribrosa deformations pre-mortem and post-mortem

**DOI:** 10.1101/2022.09.18.508448

**Authors:** Junchao Wei, Yi Hua, Bin Yang, Bo Wang, Samantha E. Schmitt, Bingrui Wang, Katie A. Lucy, Hiroshi Ishikawa, Joel S. Schuman, Matthew A. Smith, Gadi Wollstein, Ian A. Sigal

**Author notes:** Both authors should be considered co-first (Junchao Wei, Yi Hua). **Correspondence**: Ian A. Sigal, Ph.D., Laboratory of Ocular Biomechanics, Department of Ophthalmology, University of Pittsburgh School of Medicine 203 Lothrop Street, Eye and Ear Institute, Rm. 930, Pittsburgh, PA 15213, www.OcularBiomechanics.com. **Disclosures:** Junchao Wei was at the University of Pittsburgh when he contributed to this work. Junchao Wei is now at Konika-Minolta. JS Schuman receives royalties for intellectual property licensed by Massachusetts Institute of Technology to Zeiss. He is a consultant to Carl Zeiss Meditec. He is a Consultant/Advisor and is Equity owner for Opticient. He also has Intellectual property with the University of Pittsburgh. Other authors have nothing to disclose.

## Abstract

**Purpose:** Lamina cribrosa (LC) deformations caused by elevated intraocular pressure (IOP) are believed to contribute to glaucomatous neuropathy and have therefore been extensively studied, in many conditions from in-vivo to ex-vivo. We compare acute IOP-induced global and local LC deformations immediately before (pre-mortem) and after (post-mortem) sacrifice by exsanguination.

**Methods:** The optic nerve heads of three healthy monkeys 12-15 years old were imaged with spectral-domain optical coherence tomography under controlled IOP pre-mortem and post-mortem. Volume scans were acquired at baseline IOP (8-10 mmHg) and at 15, 30, and 40 mmHg IOP. A digital volume correlation technique was used to determine the IOP-induced 3D LC deformations (strains) in regions visible pre-mortem and post-mortem.

**Results:** Both conditions exhibited similar nonlinear relationships between IOP increases and LC deformations. Median effective and shear strains were, on average over all eyes and pressures, smaller post-mortem than pre-mortem, by 14% and 11%, respectively (P’s < 0.001). Locally, however, the differences in LC deformation between conditions were variable. Some regions were subjected pre-mortem to triple the strains observed post-mortem, and others suffered smaller deformations pre-mortem than post-mortem.

**Conclusions:** Increasing IOP acutely caused nonlinear LC deformations with an overall smaller effect post-mortem than pre-mortem. Locally, deformations pre-mortem and post-mortem were sometimes substantially different. We suggest that the differences may be due to weakened mechanical support from the unpressurized central retinal vessels post-mortem.

**Translational Relevance:** Additional to the important pre-mortem information, comparison with post-mortem provides a unique context essential to understand the translational relevance of all post-mortem biomechanics literature.

**Precis:** The authors compared in monkeys acute IOP-induced deformations of the lamina cribrosa pre-mortem and post-mortem. Deformation trends were similar pre-mortem and post-mortem, but deformations pre-mortem were generally smaller than those post-mortem, with substantial local variations. The differences are likely due to loss of vessel support post-mortem.

## 1. Introduction

Loss of vision in glaucoma, a leading cause of blindness worldwide, is due to damage to the retinal ganglion cell axons that transmit visual information from the light-sensitive retina to the brain.^1^ Damage to these axons is believed to initiate within the lamina cribrosa (LC), a structure within the optic nerve head through which the axons pass to exit the eye on their way to the brain.^1,2^ Elevated intraocular pressure (IOP) is a primary risk factor for glaucoma, and the only modifiable one. Nevertheless, the mechanisms by which elevated IOP contributes to neural tissue damage and vision loss are not fully understood. It is widely believed that IOP-induced deformations of the LC play a central role in the process of glaucomatous neuropathy.^2–7^

Due to technical limitations and ethical considerations, many experimental studies on LC biomechanics have been done on ocular tissues ex-vivo (after death and enucleation).^7–15^ With advances in imaging technologies, especially optical coherence tomography (OCT), there has been an increase in studies of LC biomechanics in-vivo.^4, 16–22^ Current knowledge of LC biomechanics is therefore a mix of lessons learned from a diverse mix in experimental settings, supplemented by analytic and computational modeling.^6,23–31^

It seems reasonable to suspect that there may be important differences in tissue behavior between conditions. Ocular tissues can degrade, swell or dehydrate over time, shrink during preservation, or distort during histologic processing.^32,33^ From a biomechanics perspective, enucleated globes are no longer subjected to the forces from intraocular, cerebrospinal, arterial, venous, episcleral, and orbital pressures, or from the muscles or eyelid action. There may be stress release from dissection. Some of these differences may be reduced in an ex-vivo experiment by attempting to mimic the conditions before enucleation, such as mounting the posterior pole in an inflation chamber.^34,35^ Nevertheless, these approaches are unlikely to fully reproduce the complexity of the in-vivo conditions, and therefore the potential remains that the specific conditions used for an experiment may have influenced the findings. To the best of our knowledge, the differences in LC biomechanics between in-vivo and ex-vivo conditions remain largely unknown. This precludes a proper interpretation of the findings in a given condition and an understanding of their implications on other conditions.

Our goal in this study was to compare acute IOP-induced LC deformations immediately before (pre-mortem) and after (post-mortem) death. The differences between these two, close, yet fundamentally distinct, conditions will provide information useful to understand differences between other possible experimental conditions. We adopted an experimental approach introduced by our group to image with OCT the posterior pole of healthy monkeys pre-mortem and post-mortem while at several controlled IOP levels.^4, 36–38^ We then analyzed the images to quantify acute IOP-induced LC deformations and identify any differences between pre-mortem and post-mortem responses.

## 2. Methods

Our general strategy was the following: the optic nerve heads of five eyes from three healthy adult rhesus macaque monkeys were imaged using OCT at various levels of IOP when alive (pre-mortem) and immediately after death (post-mortem). A digital volume correlation (DVC) technique was then used to find the local displacements that brought an image acquired at low IOP into coincidence with the one acquired at elevated IOP. From the displacements, we computed the 3D deformations of the LC caused by the acute increase in IOP, which we then analyzed to determine if there were differences in the LC deformations pre-mortem and post-mortem.

A note on terminology: we considered several potential terms before settling on pre-mortem and post-mortem. We discarded ex-vivo because it implies that the tissues had been removed from the body, which was not the case. We considered before and after death, but there were concerns that these did not seem “scientific”, perhaps because they are not in Latin. This was also the reason to discard “alive”, “alive condition”, and “death condition”. Other multi-word terms were considered, such as “in-situ ex-vivo”, but we preferred the simplicity of a single term. Accordingly, we opted to use pre-mortem instead of in-vivo because it afforded the symmetry of pre-mortem vs. post-mortem.

### 2.1 Animal handling and pressure set up

All procedures were approved by the University of Pittsburgh’s Institutional Animal Care and Use Committee (IACUC), and adhered to both the guidelines established in the National Institutes of Health’s Guide for the Care and Use of Laboratory Animals and the Association for Research in Vision and Ophthalmology (ARVO) statement on the use of animals in ophthalmic and vision research.

The details of animal handling and pressure setup (**Fig. 1**) have been described previously.^4,36^ Briefly, the animal was initially sedated with ketamine (20 mg/kg), diazepam (1 mg/kg), and atropine (0.04 mg/kg), and was then maintained on isoflurane (1-3%) for the remainder of the experiment. An arterial line was placed in the animal’s carotid or femoral artery, allowing for continuous recording of the intra-arterial pressure. After the arterial line was placed, the animal was put on a ventilator and was given a paralytic intravenously (vecuronium bromide, 2 mL/hour) to reduce eye movements. To record and control intracranial pressure (ICP), two small craniotomies were performed to insert a catheter (EDM lumbar catheter; Medtronic, Minneapolis, MN, USA) into the lateral ventricle and a sensor (Precision Pressure Catheter; Raumedic, Mills River, NC, USA) nearby into the parenchyma of the brain. The catheter was connected to a saline reservoir for ease of manipulating ICP (Aqualite 0.9%). IOP was controlled by inserting into the anterior chamber a 27-gauge needle, which was connected to another saline reservoir. The needle was secured to the head at the iris plane. Care was taken to ensure that it did not distort or damage the lens or cornea. To record IOP, an inline transducer was placed between the needle and the saline reservoir (TransStar, Smith Medical). M1, M2, and M3 were 12, 15, and 14 years old, respectively. Note that the same machine was used to record the ICP, IOP and arterial pressure data.

**Fig. 1.**
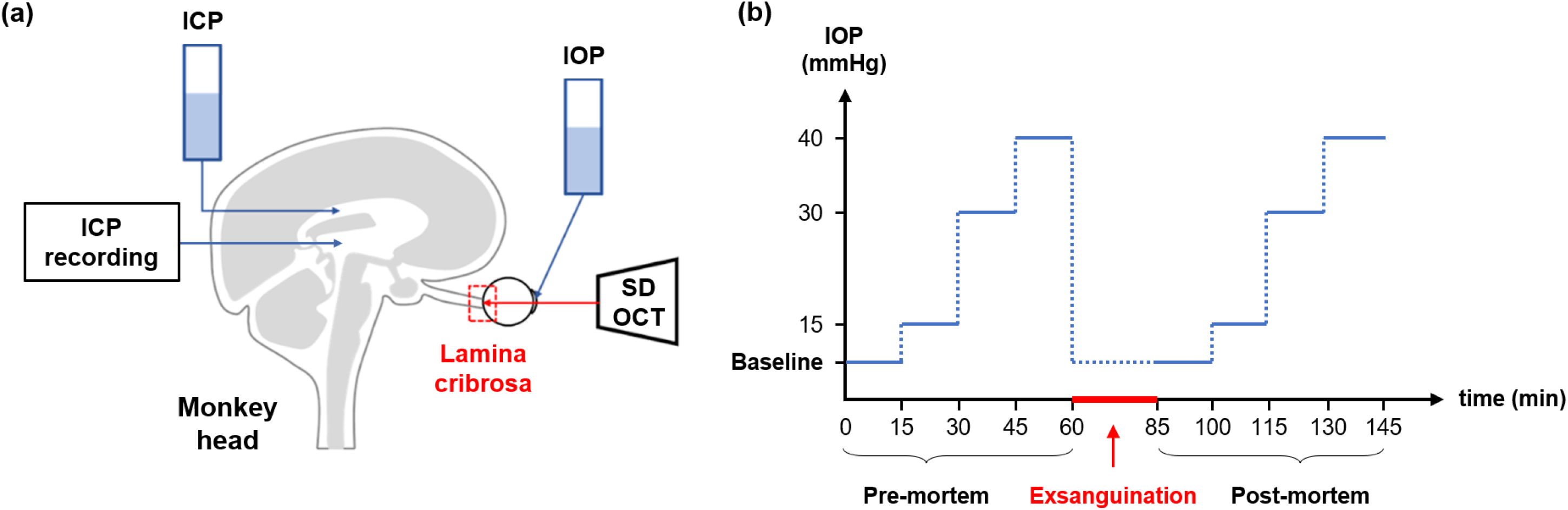
**(a)** Experimental setup modified from^36^. **(b)** Elevations of IOP in the pre-mortem and post-mortem conditions. Baseline IOP was set to 8 mmHg for M1 and M3, and 10 mmHg for M2. IOP was then raised stepwise from the baseline to 15, 30, and 40 mmHg, with each pressure step lasting about 15 minutes: 5 minutes “wait” after an IOP elevation and about 10 minutes for the scanning. For M1 and M3, we varied IOP in one eye pre-mortem and the contralateral one post-mortem. For M2, we varied IOP in the same eye pre-mortem and post-mortem. There was an interval of 25 minutes between the pre-mortem and post-mortem conditions. In both conditions, the OCT scans were first acquired at baseline IOP, and then at elevated IOPs.

#### Pre-mortem condition

For pre-mortem imaging, ICP was set to 9 mmHg, the normal ICP level in healthy animals.^39, 40^ IOP was initially set to 8 mmHg for monkey 1 (M1) and monkey 3 (M3), and 10 mmHg for monkey 2 (M2). IOP was then raised stepwise from the baseline pressure to 15, 30, and 40 mmHg, with each pressure step lasting approximately 15 minutes. IOP, ICP, and blood pressure were measured and continuously recorded digitally at a rate of 100 Hz (MPR 1 DATALOGGER; Raumedic).

#### Exsanguination and post-mortem condition

Before exsanguination, 3000 to 5000 units of heparin were given intravenously and isoflurane was increased to the upper end of the specified range to ensure deep anesthesia. The intravenous saline drip was also left open during the exsanguination to assist flow. Suction was then added to the arterial line with a vacuum pressure of 584 mmHg. The valve was periodically closed to obtain a blood pressure reading, and the suction was then continued. Animal death was defined as the moment when there was no longer a heartbeat or a blood pressure reading and no pulsation of blood vessels on the OCT viewer. In the post-mortem condition, we controlled and measured IOP only.

### 2.2 Image acquisition

The animal was placed in a prone position during OCT imaging. The eyes were kept open using a wire speculum. Tropicamide drops (0.5%, Bausch & Lomb, Rochester, NY, USA) were used to dilate the pupils. The cornea was fitted with a rigid gas-permeable contact lens (Boston EO, Boston, MA, USA) to improve image quality and kept hydrated throughout the experiment with artificial tears.

Spectral-domain OCT (Leica, Chicago, IL, USA), with a broadband superluminescent diode light source (Superlum, Dublin, Ireland; λ = 870 nm, Δλ = 200 nm), was used to acquire 3D volume scans (3 mm × 3 mm × 1.92 mm with 400 × 400 × 1024 voxels sampling) of the optic nerve head region, focused on the LC. The voxels of the OCT volumes were therefore 7.5 μm × 7.5 μm × 1.875 μm. In both the pre-mortem and post-mortem conditions, these scans were first acquired at baseline IOP and then at progressively elevated IOPs. To reduce potential viscoelastic effects, all scans were acquired at least 5 minutes after a given IOP change.^35^ Multiple scans were collected at each IOP level, with various focal depth settings. The scans with the best LC microstructure visualization were selected for analysis.

### 2.3 Image analysis

#### Motion artifact correction

Motion artifacts in the OCT scans result from breathing and heart rate, as well as the environmental vibration of the OCT device and surgical table. To remove the motion artifacts, a traditional method is to apply a non-rigid affine transformation to the scans.^41^ However, this method is aggressive and may erroneously remove the true deformations. To avoid this problem, in this work, we utilized a technique that first registered the scans sequentially based on the optic nerve head structures, and then corrected motion artifacts using a causal low-pass filter during the registration.^42^ A comparison of the OCT scans before and after the correction of motion artifacts is shown in **Fig. 2**.

**Fig. 2.**
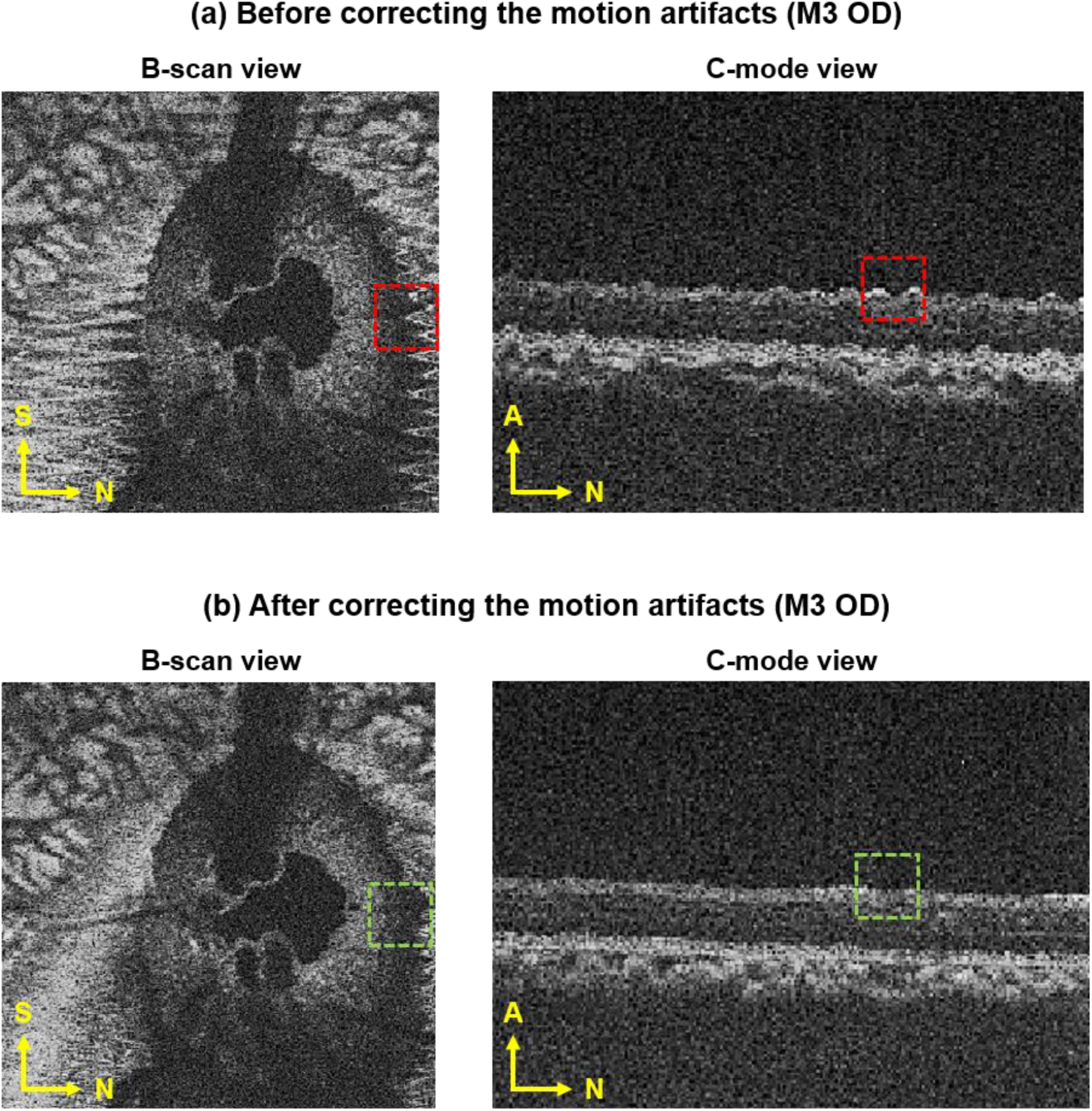
A comparison of OCT scans **(a)** before and **(b)** after the correction of motion artifacts. To remove the motion artifacts in OCT scans (red dotted boxes), we first registered the scans sequentially based on the optic nerve head structures, and then corrected the artifacts using a causal low-pass filter during the registration. Panel (b) shows the efficiency of our method to remove the motion artifacts (green dotted boxes). S – Superior, N – Nasal, A – Anterior.

#### Noise reduction

The speckle-like noise in the OCT scans is an inherent artifact in coherence imaging, which can reduce the image contrast and affect both the axial and lateral resolutions.^43^ We applied a 3D median filter on the OCT volume scans for noise reduction.^44^ A nonlinear sigmoid transfer function was then used to further improve image contrast.^45^ A comparison of the OCT scans before and after the noise reduction is shown in **Fig. 3**.

**Fig. 3.**
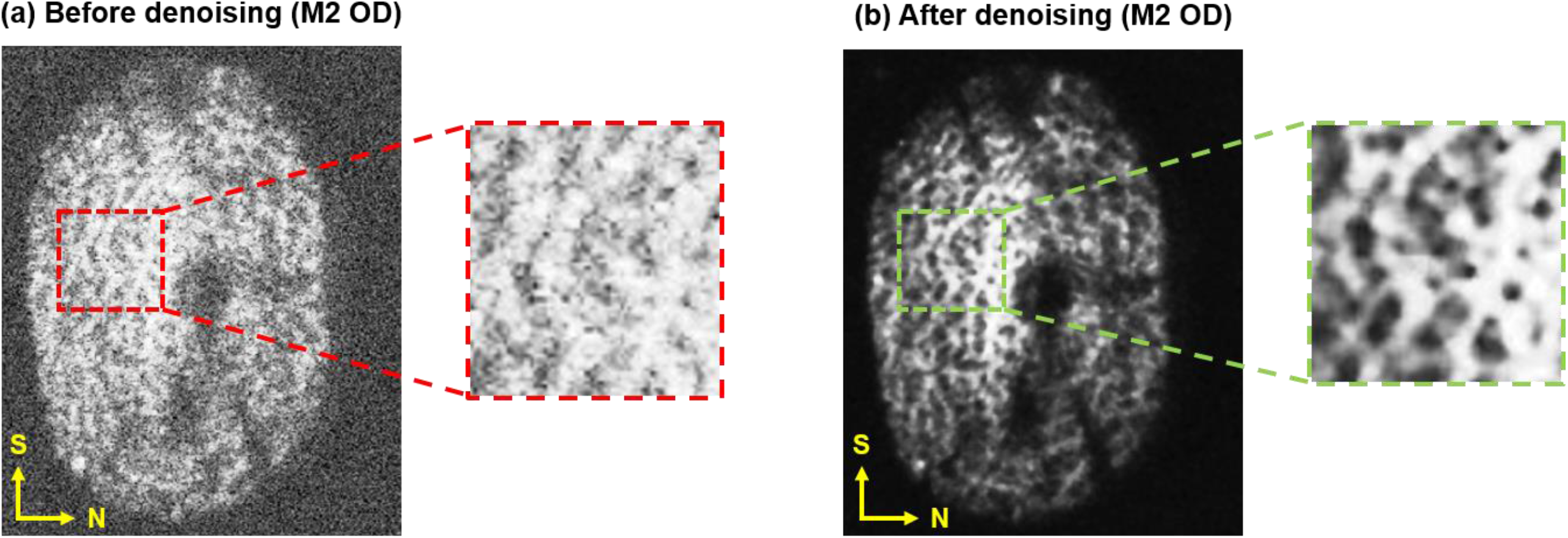
A comparison of OCT scans **(a)** before and **(b)** after denoising. We applied a 3D median filter on the OCT volume scans for noise reduction. A nonlinear sigmoid transfer function was then used to further improve the image contrast. Our method was efficient in reducing the speckle-like noise in OCT scans. After denoising, the LC trabecular beams and pores (close-ups) were more visible than those before denoising. S – Superior, N – Nasal.

#### Digital volume correlation

To determine the IOP-induced LC deformations, we used a DVC analysis between multiple OCT scans.^42^ For simplicity, the scans were treated as 3D volumetric images. The basic concept of DVC is to establish a mapping between an undeformed volume at baseline IOP (8 mmHg for M1 and M3, 10 mmHg for M2) and a deformed one at elevated IOP. The undeformed volume was treated as a baseline volume, then used to compare with each deformed volume. The result of this mapping was a transformation function between two volumes. The transformation function was evaluated by minimizing the error of zero-normalized cross-correlations, which is a numerical metric for finding a similarity.^46^ We applied the Nelder-Mead method to minimize the error of zero-normalized cross-correlations.^47^ The minimization of the correlation values indicates the best matching point in the search region of the deformed volume, and from the matched points, the displacement vector can be calculated. We computed the Green-Lagrange strain tensor from the displacement field at the matched points. The effective strain (a measure of the total deformation including stretch and compression) and the maximum shear strain (a measure of tearing and bending deformations) can be extracted as

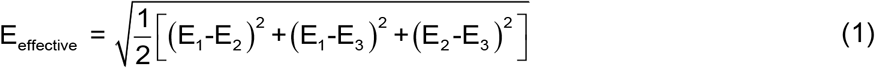

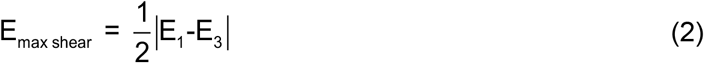

where E_1_, E_2_, and E_3_ are the principal components of the Green-Lagrange strain tensor. For simplicity, the maximum shear strain is hereafter referred to as shear strain.

#### Masking

As we have previously reported, exsanguination increases the visibility of the LC.^36^ To make a fair comparison of the IOP-induced LC strains between pre-mortem and post-mortem conditions, we had to compare only the regions visible in both conditions. For this, the blood vessel shadows in the pre-mortem scans were manually selected and used as a mask on the post-mortem scans (**Fig. 4**). Other low-quality correlations were identified using a statistical z-score and removed to ensure that unreliable displacements were not included for computing deformations or considered in the comparison. For monkeys 1 and 3, the image quality post-mortem was poor, which substantially reduced the regions of reliable analysis. Thus, for these animals, the comparisons between pre-mortem and post-mortem conditions were made with the contralateral eye. This required carefully registering and mirroring the images of the contralateral eye.

**Fig. 4.**
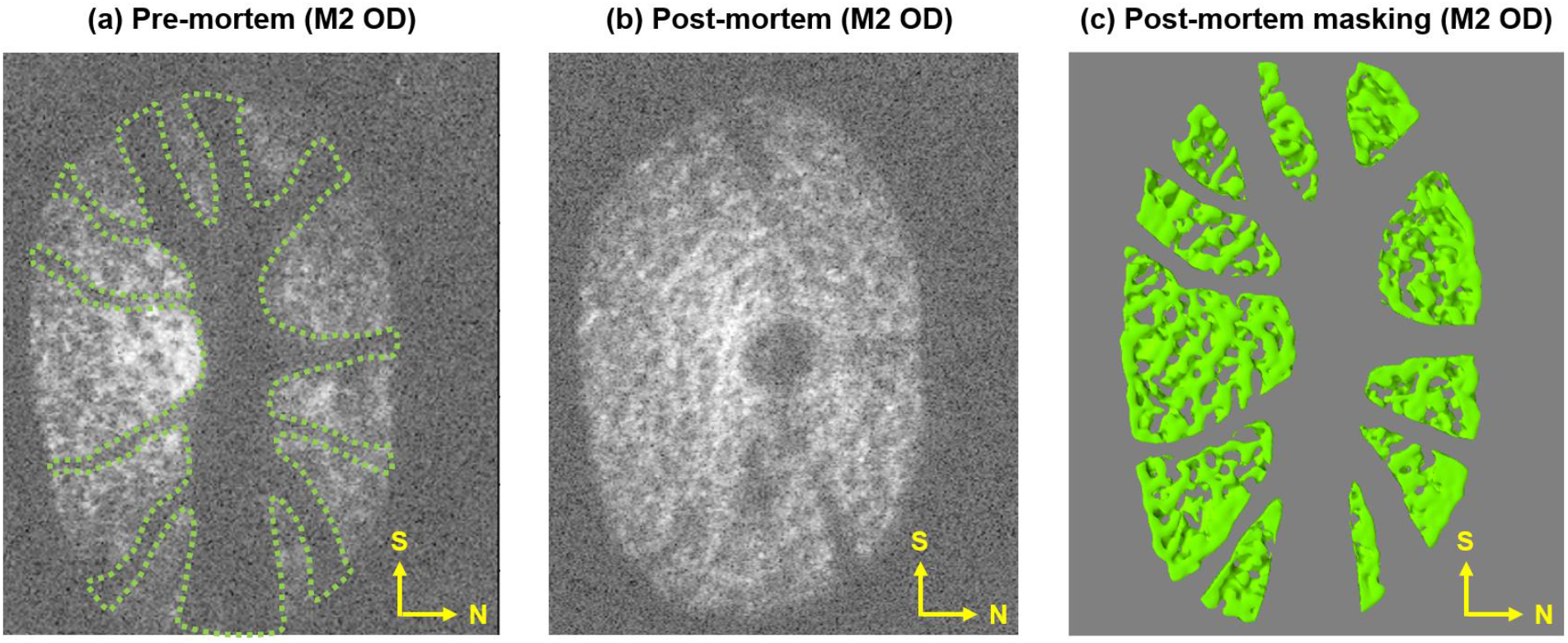
Demonstration of shadow masking. **(a)** C-mode view of the LC at baseline IOP in the pre-mortem condition. The outline of blood vessel shadows was manually delineated and used to identify the region for comparison. **(b)** C-mode view of the LC at baseline IOP in the post-mortem condition. **(c)** The same scan in the post-mortem condition masked using the outline from the scan in the pre-mortem condition. S – Superior, N – Nasal.

### 2.4 Statistical analysis

We determined whether the IOP-induced LC strains were significantly different between pre-mortem and post-mortem conditions using linear mixed-effects models. As fixed variable we used the condition pre-mortem or post-mortem. The random variables were factors that may affect the sampling population, such as the level of IOP. Our analysis included voxels from the whole lamina for both the pre-mortem and post-mortem OCT volume scans. The statistical analysis was based on points tracked within the visible LCs. Sample sizes were 2,249 for M1, 1,731 for M2, and 2,221 for M3. We removed the spatial autocorrelation by specifying an exponential spatial correlation structure using the cartesian distance between voxels within each scan volume.^48^ We used alpha = 0.001 to establish significance. Statistical analysis was done with R v3.2.2 (R Foundation for Statistical Computing).

## 3. Results

In both the pre-mortem and post-mortem conditions, our image registration technique performed well to match low and high IOP images (**Fig. 5**). Before the registration, the differences in images cannot be removed by simple translation or rotation. Conversely, after the registration, there was an excellent match between images with the overlapping regions of lamina beams and pores. The minor differences in images are due to the noise and intensity variations that result in the slightly greenish or reddish regions despite the coincidence of the LC features.

**Fig. 5.**
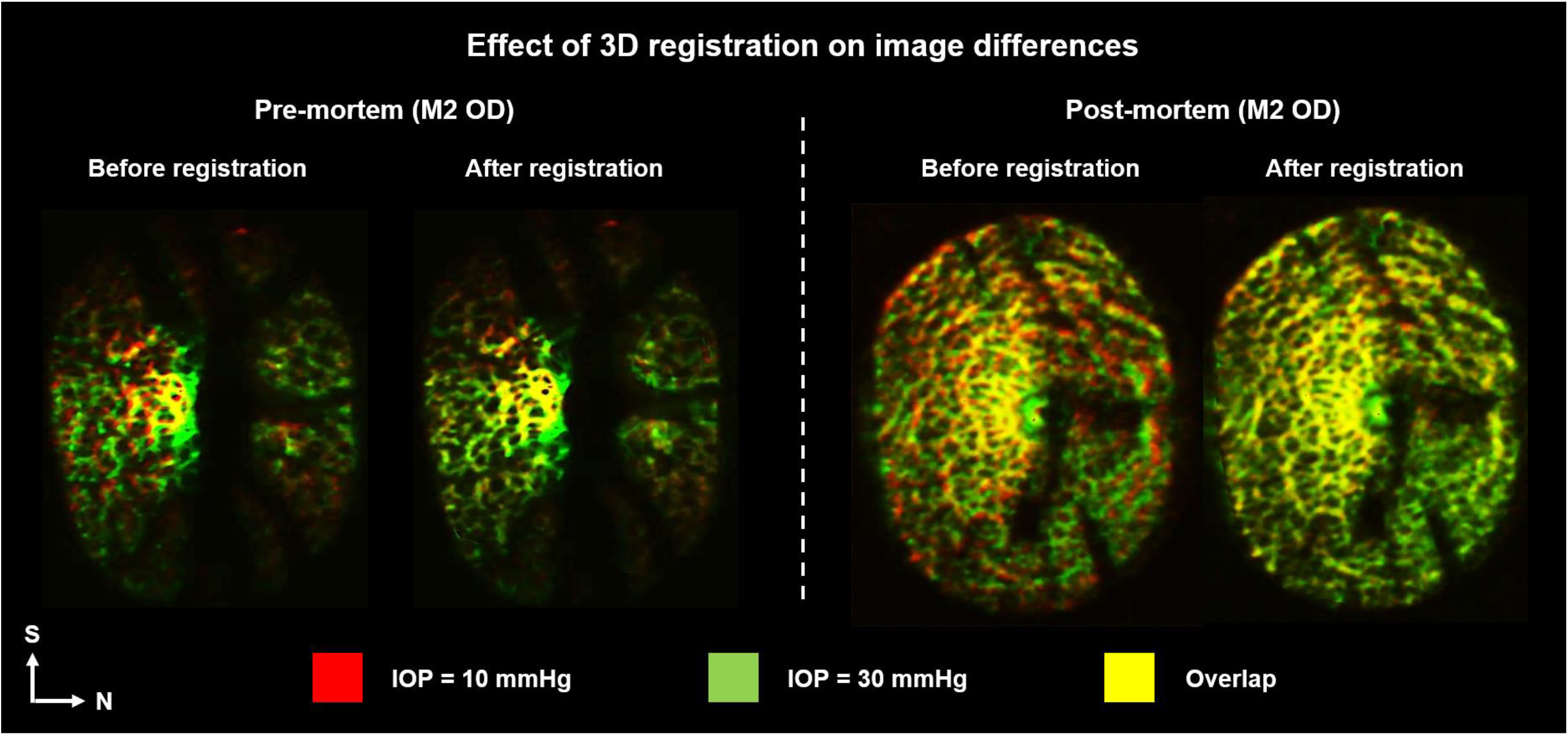
Demonstration of the 3D image registration technique in the pre-mortem (left) and post-mortem (right) conditions. Shown are OCT C-mode view acquired at IOPs of 10 (red) and 30 mmHg (green) to visualize the differences and overlap (yellow). The differences before registration cannot be removed by simple translation or rotation. The 3D registration produced excellent coincidence between the images, without concentrations. The largest differences were due to noise and intensity variations that result in slightly greenish or reddish regions despite the coincidence of the LC features. S – Superior, N – Nasal.

The LC strains increased with IOP in both the pre-mortem and post-mortem conditions (**Figs. 6–8**). IOP-induced LC strains were highly focal and concentrated in regions as small as a few pores. Maps of the image differences before and after registration demonstrate that this was not a registration artifact (**Fig. 5**). Overall, IOP-induced LC strains were smaller post-mortem than pre-mortem. Locally, some regions were subjected pre-mortem to triple the strains observed post-mortem, whereas a few locations had larger strains post-mortem than pre-mortem (**Fig. 9**).

**Fig. 6.**
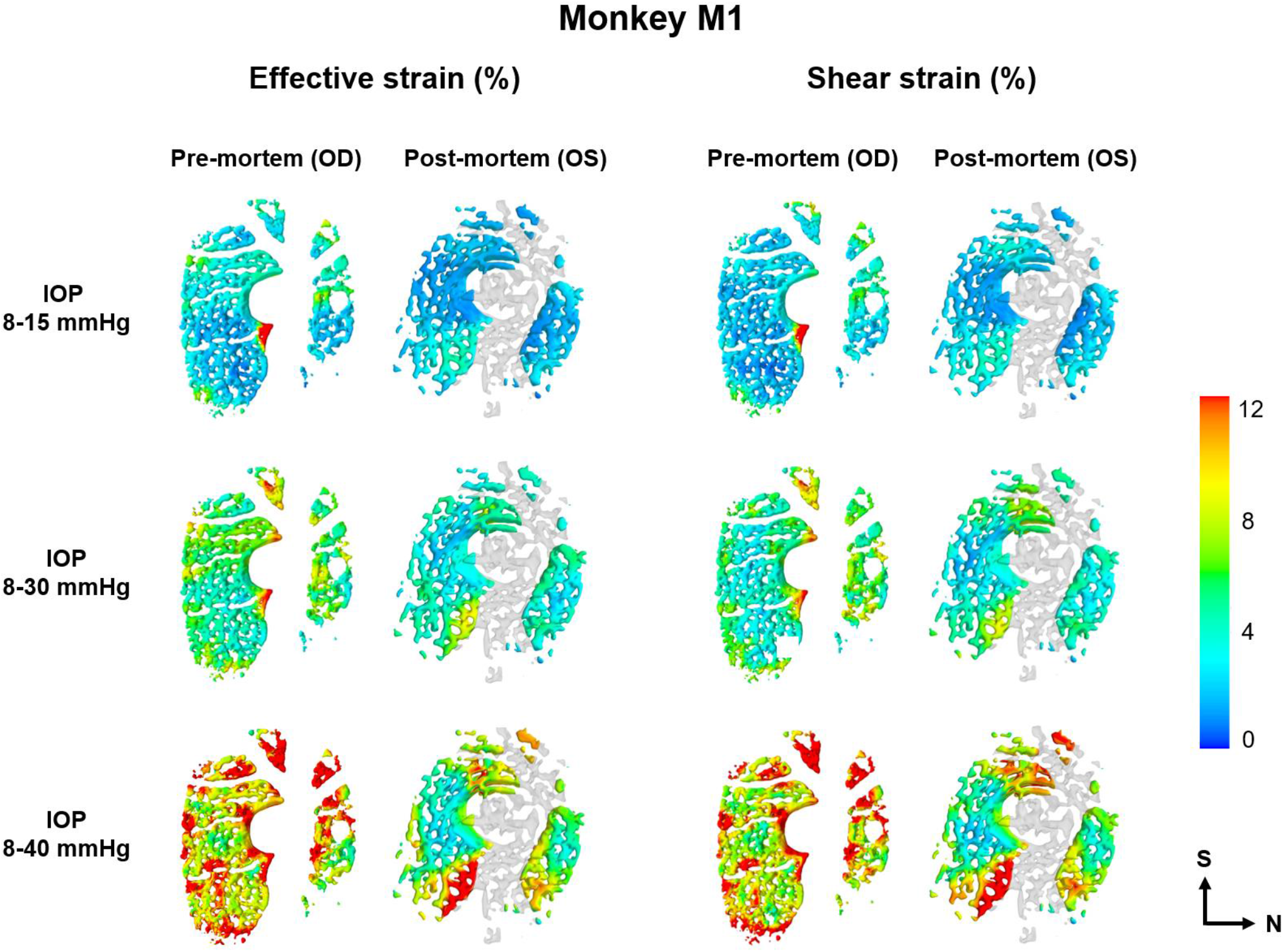
Contour plots of the IOP-induced LC strains of monkey **M1** in the pre-mortem and post-mortem conditions. Left two columns: the effective strain in the pre-mortem and post-mortem conditions, right two columns: the shear strain in the pre-mortem and post-mortem conditions. Rows 1-3 correspond to the three levels of IOP elevation (from low to high). The distribution of the effective strain was similar to that of the shear strain, which were highly focal and concentrated in regions as small as a few pores. IOP-induced LC strains were overall smaller post-mortem than pre-mortem, but locally, a few locations had larger strains post-mortem than pre-mortem. OS images were flipped for ease of comparison. S – Superior, N – Nasal.

**Fig. 7.**
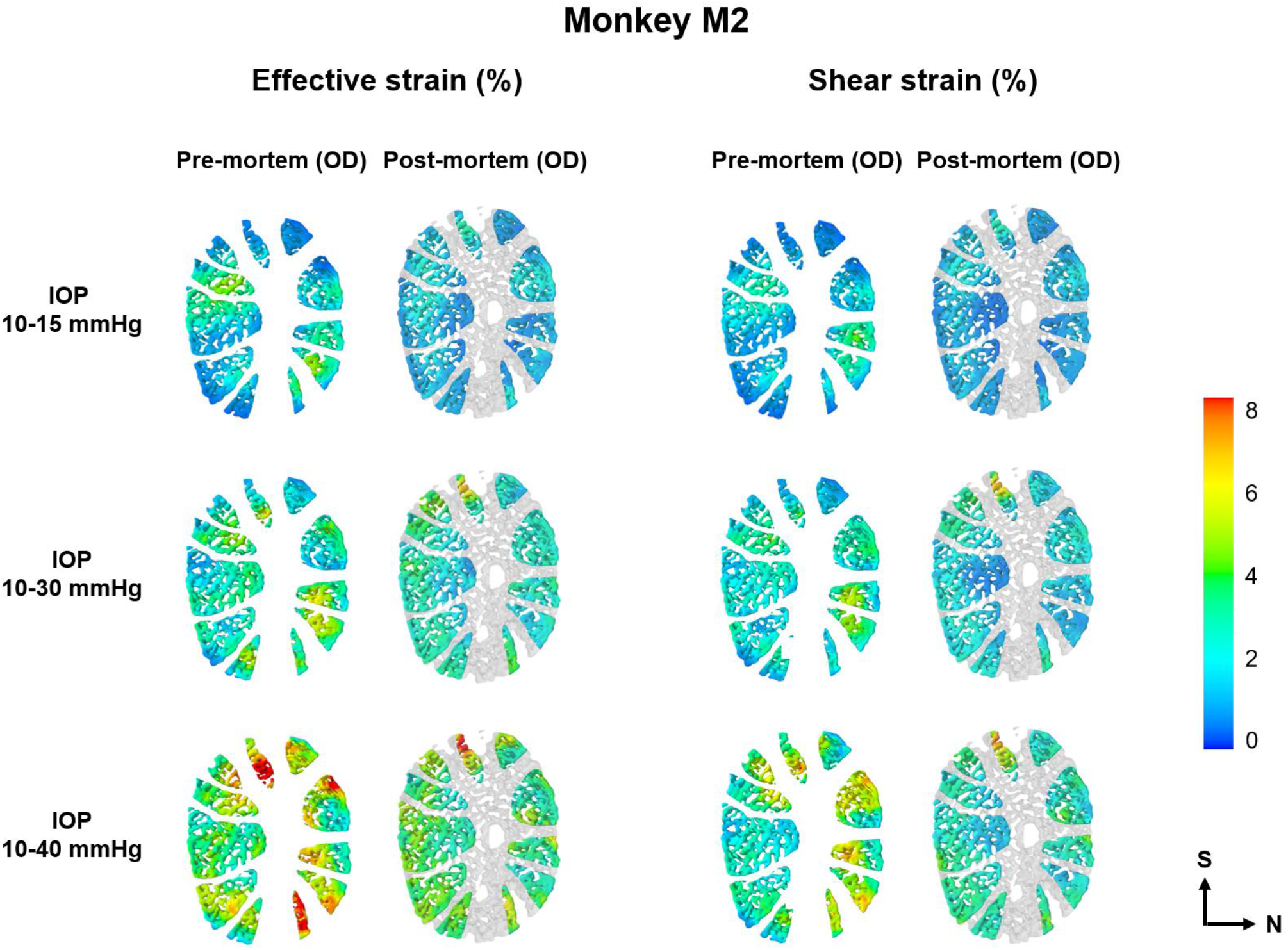
Contour plots of the IOP-induced LC strains of monkey **M2** in the pre-mortem and post-mortem conditions. Left two columns: the effective strain in the pre-mortem and post-mortem conditions, right two columns: the shear strain in the pre-mortem and post-mortem conditions. Rows 1-3 correspond to the three levels of IOP elevation (from low to high). Similar to the observations of M1 (Fig. 6), IOP-induced LC strains were overall smaller post-mortem than pre-mortem, but locally, a few locations had larger strains post-mortem than pre-mortem. Note that the color scale of these plots is different from that of Figs. 6 and 8. The range was selected to help discern details of the patterns. S – Superior, N – Nasal.

**Fig. 8.**
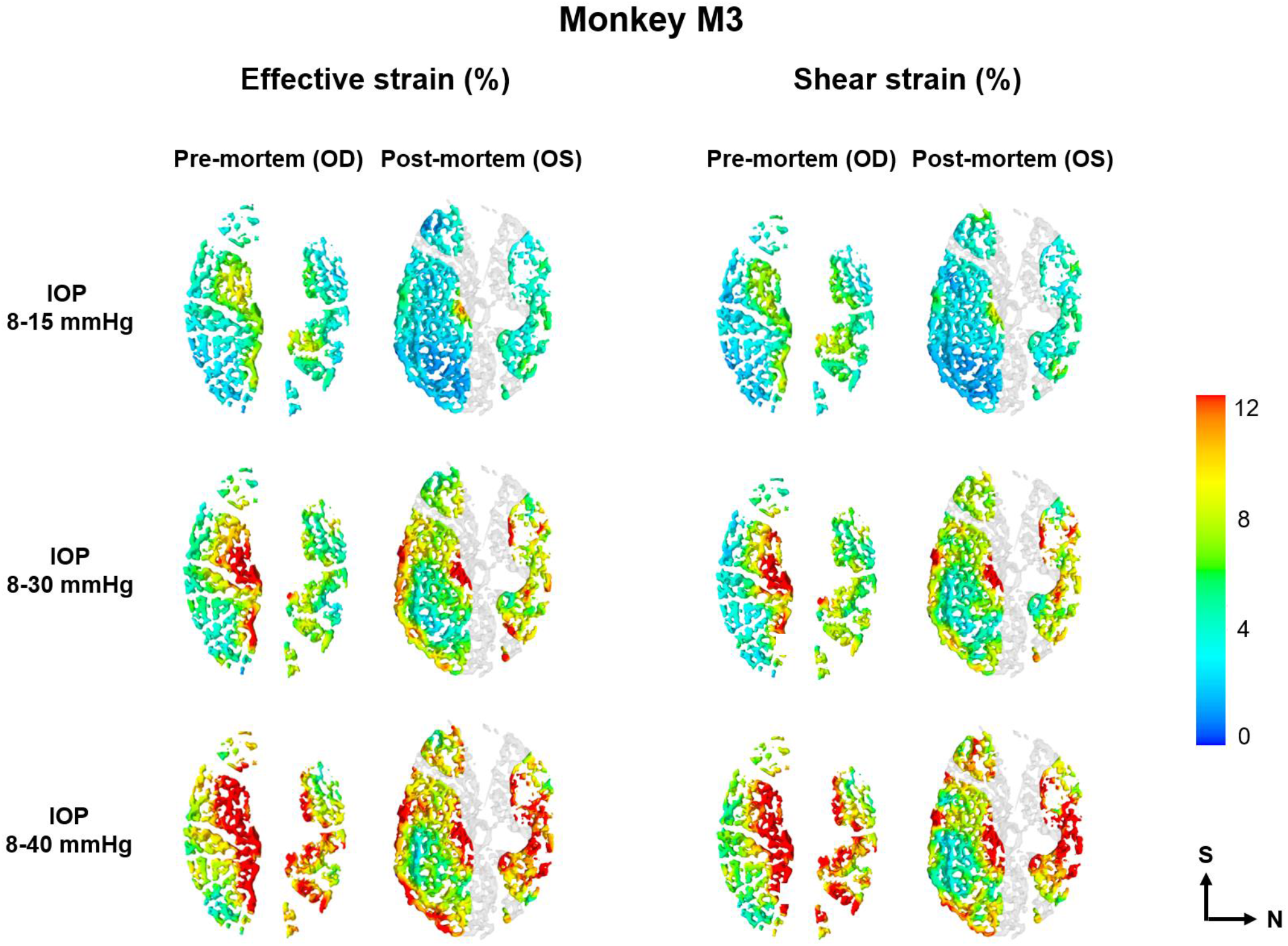
Contour plots of the IOP-induced LC strains of monkey **M3** in the pre-mortem and post-mortem conditions. Left two columns: the effective strain in the pre-mortem and post-mortem conditions, right two columns: the shear strain in the pre-mortem and post-mortem conditions. Rows 13 correspond to the three levels of IOP elevation (from low to high). Similar to the observations of M1 (Fig. 6), the IOP-induced LC strains were overall smaller post-mortem than pre-mortem, but locally, a few locations had larger strains post-mortem than pre-mortem. S – Superior, N – Nasal.

**Fig. 10** shows a quantitative comparison of the IOP-induced LC strains between pre-mortem and post-mortem conditions. On average, over the three monkeys, the effective and shear strains decreased by 14.4% and 11.0%, respectively, in the post-mortem condition relative to those in the pre-mortem condition. The largest decreases were in M3 when IOP increased from 8 to 15 mmHg, in which the effective and shear strains decreased by 23.3% and 16.6%, respectively (P < 0.001). Compared with the pre-mortem condition, IOP-induced LC strains in the post-mortem condition were less variable.

**Fig. 9.**
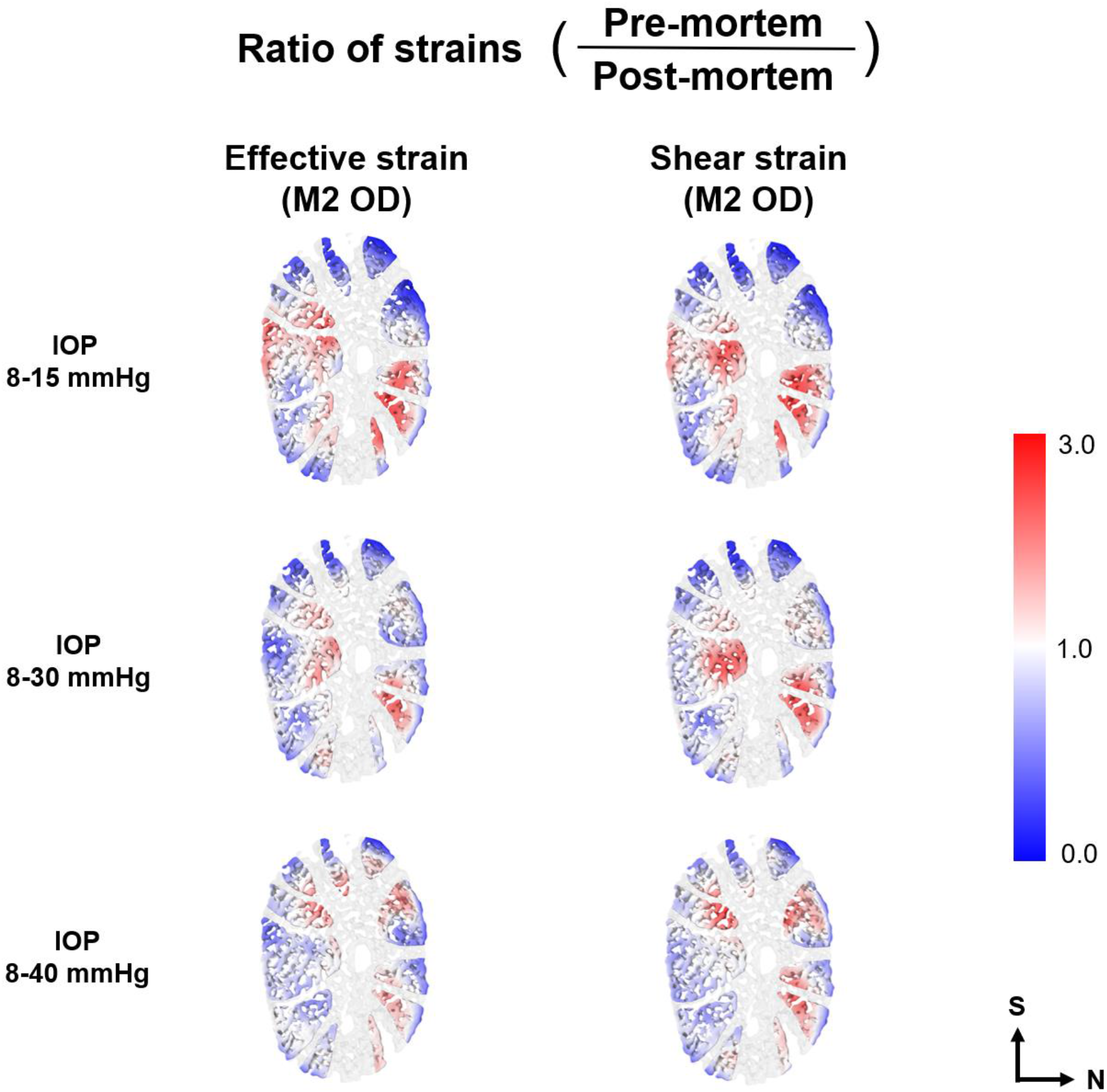
Contour plots of the ratio of IOP-induced LC strains between pre-mortem and post-mortem conditions (M2 OD). Left column: the effective strain, right column: the shear strain. Rows 1-3 correspond to the three levels of IOP elevation (from low to high). Larger strains in the pre-mortem condition are shown in red and smaller strains in blue. Some regions were subjected pre-mortem to triple the strains than post-mortem (red). A few locations had larger strains post-mortem than pre-mortem (blue). S – Superior, N – Nasal.

**Fig. 10.**
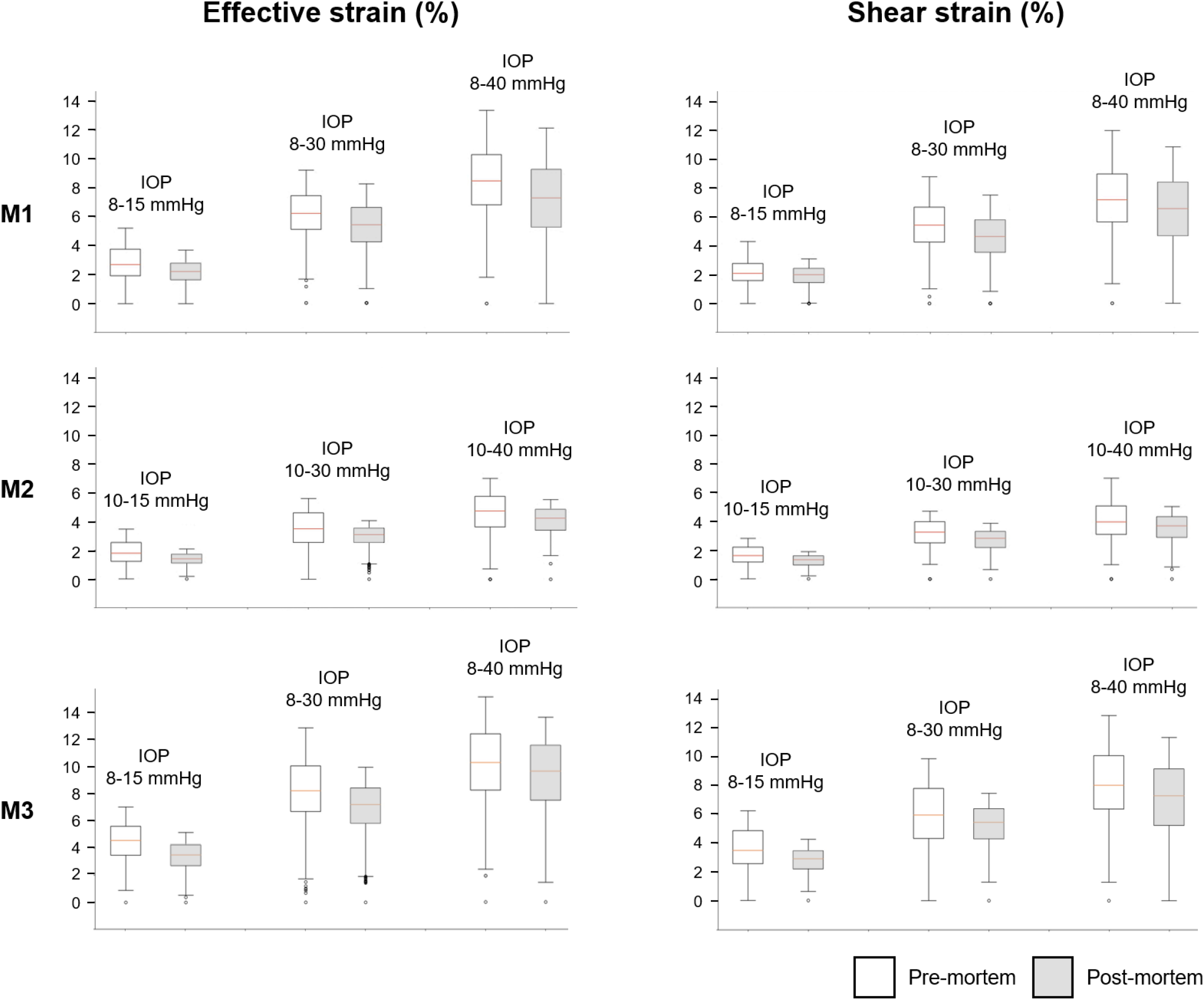
Boxplots of IOP-induced LC strains in the pre-mortem and post-mortem conditions. Left column: the effective strain, right column: the shear strain. Rows 1-3 correspond to the strain measurements in M1-M3. Overall, IOP-induced LC strains post-mortem were smaller and less variable than pre-mortem. On average, the effective and shear strains decreased by 14.4% and 11.0%, respectively, in the post-mortem condition relative to those in the pre-mortem condition. The largest decreases were in M3 when IOP increased from 8 to 15 mmHg, in which the effective and shear strains decreased by 23.3% and 16.6%, respectively (P < 0.001).

The LC median strains increased nonlinearly with IOP in both the pre-mortem and post-mortem conditions (**Fig. 11**). For example, when IOP increased from 15 to 30 mmHg, the effective strain increased by an average of 101.6% and 123.1%, respectively, in the pre-mortem and post-mortem conditions. In comparison, when IOP increased from 30 to 40 mmHg, the effective strain only increased by an average of 32.5% and 35.4%, respectively, in both conditions.

**Fig. 11.**
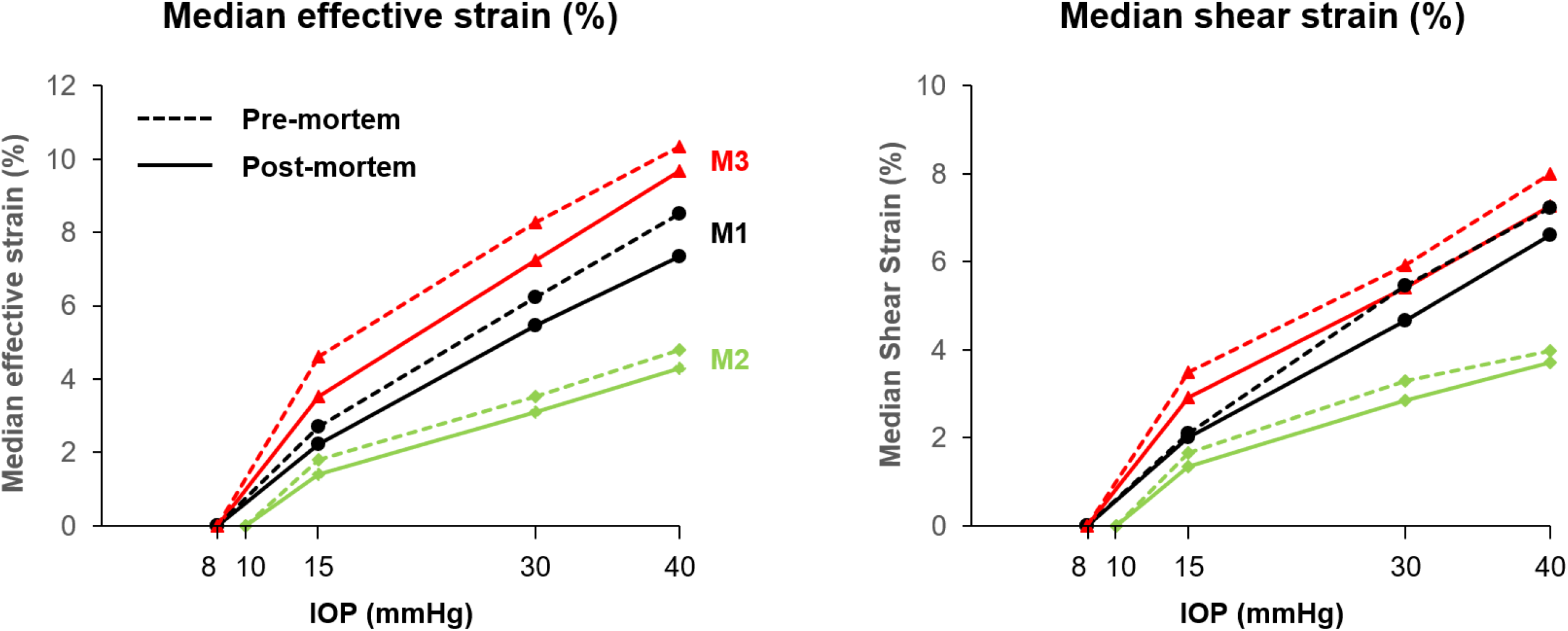
Comparison of IOP-induced LC median strains in the pre-mortem (dashed line) and post-mortem (solid line) conditions of M1M3. Left: the median effective strain; Right: the median shear strain. For all the three monkeys, the median effective and shear strains increased nonlinearly with IOP in both the pre-mortem and post-mortem conditions. Although the values were different, the trends were similar pre-mortem and post-mortem.

## 4. Discussion

Our goal was to compare the acute IOP-induced 3D LC deformations immediately before and after death. Overall, IOP-induced LC deformations were smaller post-mortem than pre-mortem, but the trends in the nonlinear relationships with IOP were similar. Locally, deformations pre-mortem and post-mortem were sometimes substantially different, with some regions suffering much smaller and some larger post-mortem deformations than pre-mortem.

This study provides important information that we have not found elsewhere on the differences between the effects of IOP on the LC pre-mortem and post-mortem. We postulate that the decreased IOP-induced LC strains post-mortem can be explained by weakened mechanical support from the central retinal vessel post-mortem due to the loss of hydraulic stiffness after exsanguination (**Fig. 12**). Without the support from the vessel, the central LC displaces posteriorly, even at normal IOP. From this condition, the increases in IOP cause deformations that are smaller post-mortem in the central LC and larger in the peripheral LC. This interpretation is supported by a direct visualization of OCTs overlaid on each other (**Fig. 13**).

**Fig. 12.**
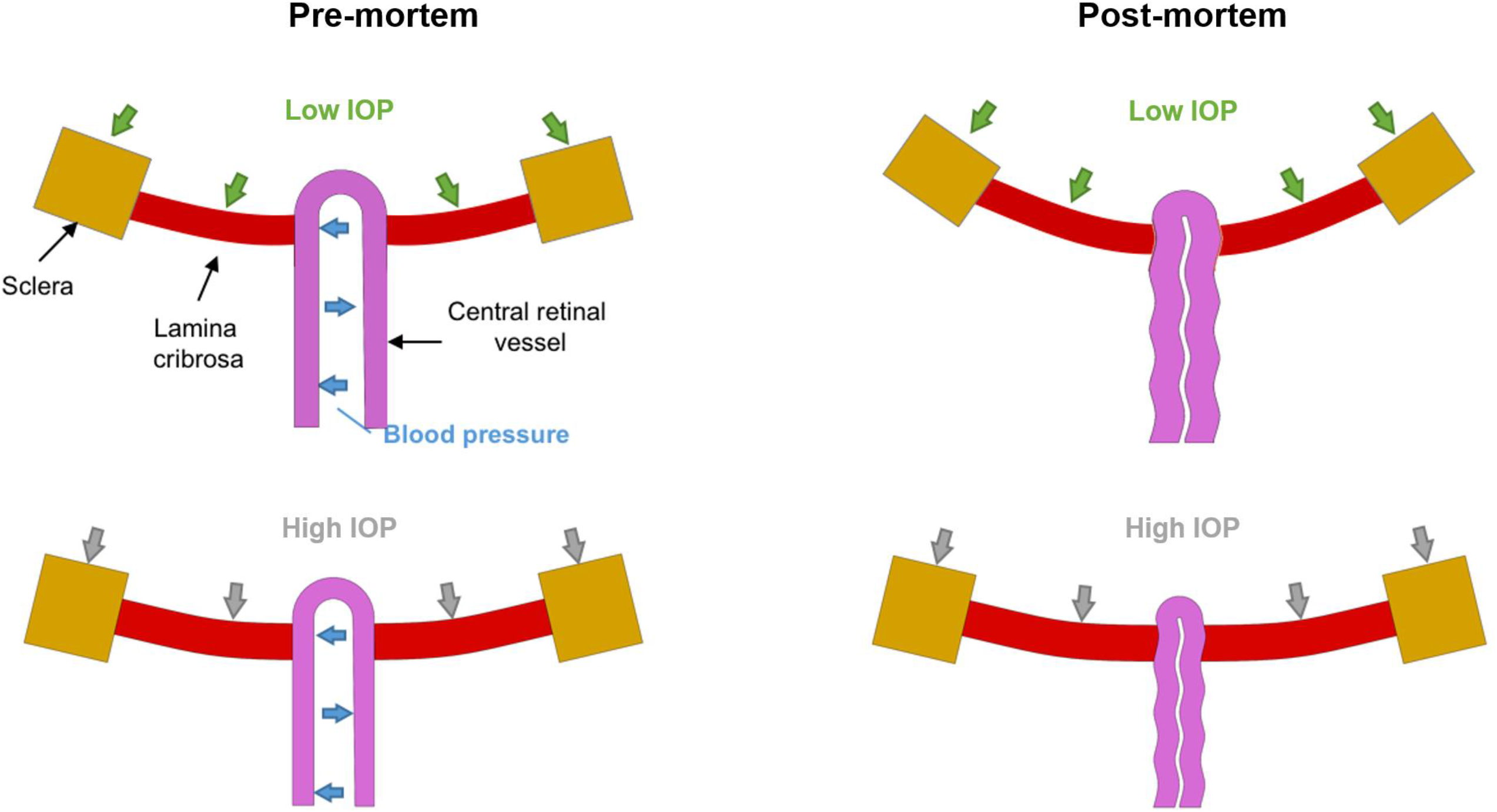
Schematic representation of how the central retinal vessel influences the IOP-induced LC deformations in the pre-mortem and post-mortem conditions. In this framework, in the pre-mortem condition, the central retinal vessel acts like a tent pole to support the central region of the LC.^28,67^ As IOP increases, the direct effects of IOP “push” the LC posteriorly, and the indirect effects deform the sclera, expanding the canal, which in turn “pull” the LC taut from the sides.^68^ After exsanguination, even at normal IOP, the vessel collapses due to the loss of blood pressure.^36^ Without the support of vessel, the central region of the LC in the post-mortem condition also moves posteriorly relative to the pre-mortem condition. As IOP increases, further posterior displacement of the LC is limited, resulting in the decreased LC strains in the post-mortem condition.

**Fig. 13.**
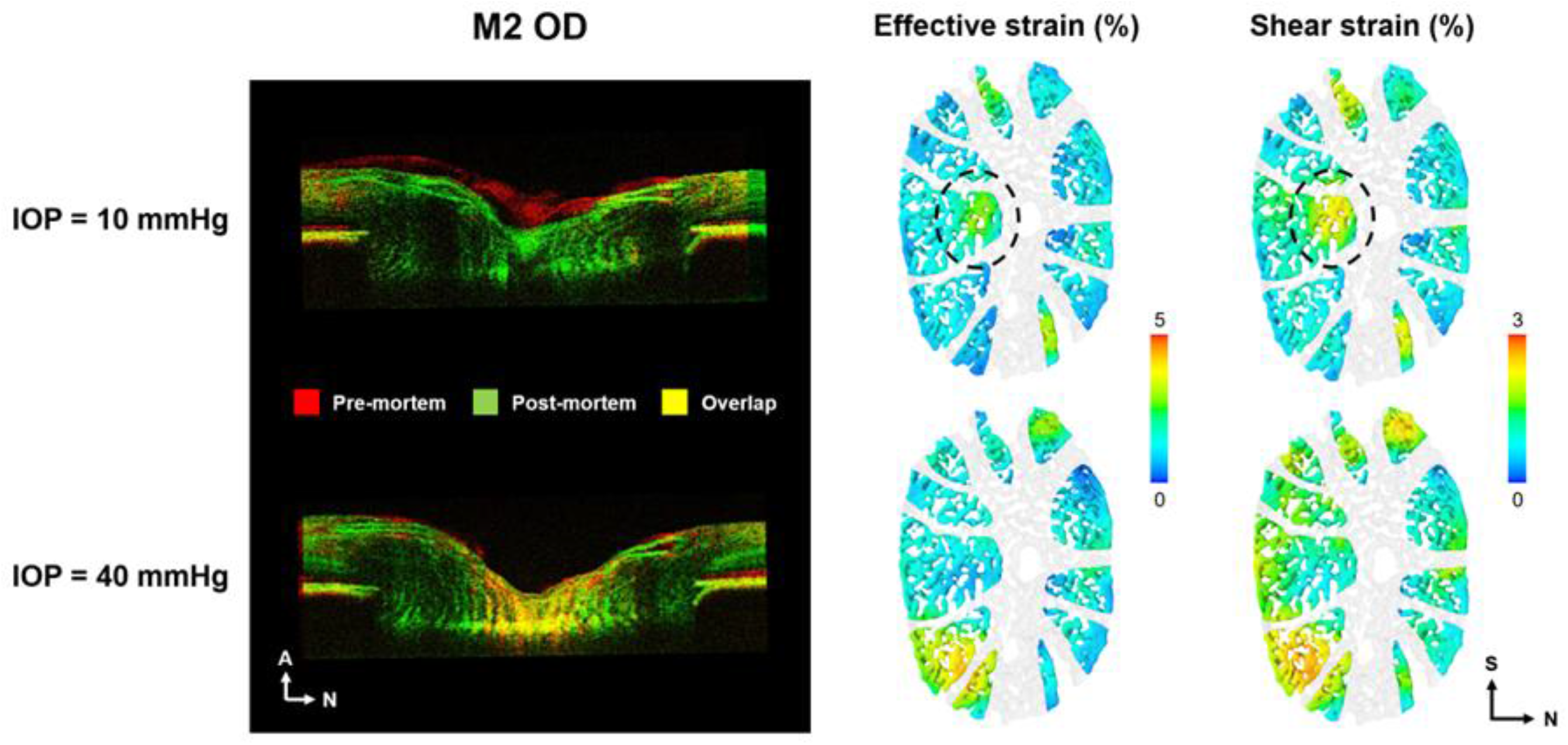
Comparison of the optic nerve head between pre-mortem and post-mortem conditions. Shown on the **left** are OCT B-scans for M2 OD acquired at IOPs of 10 (top) and 40 mmHg (bottom). The pre-mortem scan is shown in red, and the post-mortem scan is shown in green, with yellow representing overlap. Shown on the **right** are the effective and shear strains of the LC calculated by DVC between pre- and post-mortem conditions. At baseline IOP (10 mmHg), the central LC in the post-mortem condition was displaced more posteriorly relative to the pre-mortem condition. This is consistent with the DVC measurements showing larger strains in the central region of the LC (as indicated by the dashed circle). A potential explanation is that, in the pre-mortem condition, the central retinal vessels are under hydraulic stiffening from blood pressure and thus provide support to the adjacent tissues. After exsanguination, the vessels collapse due to the loss of blood pressure, resulting in the tissues being displaced posteriorly. It is also interesting that the large displacements were accompanied by LC strains comparable to those caused by fairly large changes in IOP. As IOP increases to 40 mmHg (bottom), further posterior displacement of the LC is limited in the post-mortem condition, resulting in smaller LC strains post-mortem than pre-mortem. N – Nasal, A – Anterior, S – Superior.

Our finding of the decreased IOP-induced LC strains post-mortem than pre-mortem is consistent with recent measurements in human, monkey, and porcine eyes. For example, pre-mortem studies on human^21^ and monkey^22^ eyes reported that strains in the LC could reach 10% to 20% as IOP increased by a range of 10-23 mmHg. In comparison, post-mortem studies on human^12,13,15^ and porcine^14^ eyes generally measured the LC strains less than 10% as IOP increased by a range of 10-40 mmHg. The differences in the IOP-induced LC strains between pre-mortem and post-mortem conditions are likely smaller than those between pre-mortem and ex vivo studies done after the eye has been enucleated and dissected. Enucleated eyes have more differences with the pre-mortem condition that might affect the regional mechanical response, including loss of orbital support, changes in choroidal volume,^49^ and longer times between death and testing, sometimes of multiple days.^50^ Thus, our measurements also provide a lower boundary estimate of the combined effects of sacrifice and enucleation.

The observed nonlinear relationship between IOP and LC median strains in both conditions is likely primarily determined by tissue property nonlinearities, which are, in turn, determined in large part by collagen fiber undulations, or crimp.^51–54^ Using polarized light microscopy, we have reported microstructural crimp of collagen fibers in the LC and sclera at normal IOPs.^52^ The crimp gradually vanished as IOP increased, contributing to the local stiffening of the tissue.^51^ This process is called collagen fiber recruitment, which is well recognized in other tissues, like tendons and ligaments.^55–57^ It is important to acknowledge that, while crimp plays a central role in soft tissue mechanics, it is not the only factor affecting the mechanical properties. Several other factors also affect tissue stiffness and the response to load, including, but not limited to, fiber type, composition, thickness, alignment and slip conditions, proteoglycan and elastin content and distribution, the amount and type of cross-linking between fibers, and the presence and characteristics of blood vessels within.^55, 58–63^

A key strength of our study is that the IOP-induced LC strains were measured in 3D by DVC pre-mortem and post-mortem. The LC is intrinsically a 3D structure and capturing its deformations in 3D renders more robust measurements and a better representation of the actual deformations than a 2D approximation. To improve the accuracy of our DVC measurements, we corrected the motion artifacts and reduced the speckle-like noise in OCT scans before image registration. We also examined the correlation coefficients after the DVC measurements (**Fig. 14**). We then verified our DVC measurements by recovering the undeformed scans from applying the calculated displacement vectors to the deformed ones. Measuring the LC deformations at multiple levels of IOP allowed us to track the continuous deformation of the LC and to capture the nonlinear relationship between IOP and LC strains. The combination of OCT imaging and DVC analysis can be further used to track changes, acute or chronic. This could prove powerful to characterize the patterns of damage and causes underlying susceptibility to pathology in eyes with chronic elevated IOP.

**Fig. 14.**
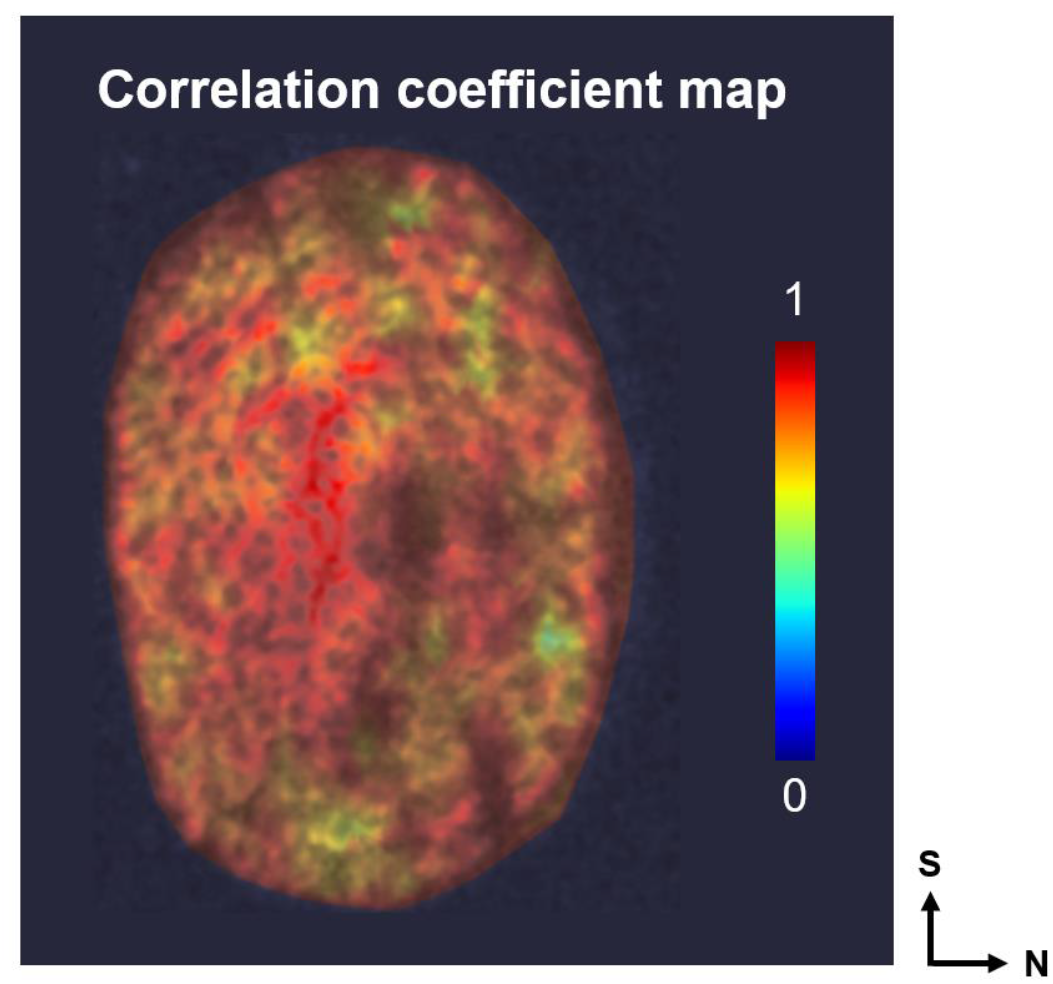
An example of correlation coefficient map overlaid on the OCT image (M2 OD post-mortem). The high correlations indicate a robust DVC analysis. Note that the correlation coefficients are discrete. We interpolated the coefficient values to generate the contour plot.

Another strength of our study was the correction for spatial autocorrelation done in the statistical analysis. Autocorrelations that remain uncorrected or not accounted for can spuriously increase the confidence in results. We would like to give credit to Fazio et al. for bringing to our attention this important yet extremely uncommon step in the analysis.^48^

It is also important to consider the limitations of our methods when interpreting the results. First, the cornea was fitted with a contact lens to improve image quality. Although the benefits of using the contact lens for imaging make it worthwhile, it may have affected the response of the globe to IOP and the biomechanics of the posterior pole.^64^ However, the lens was present in both pre-mortem and post-mortem conditions, so the effects on the comparative results observed in this work are expected to be minor, and unlikely to affect the conclusions overall.

Second, due to the substantial shadowing from the vasculature in the pre-mortem scans, the differences in the IOP-induced LC strains between pre-mortem and post-mortem conditions were determined by comparing only measurements from the same visible regions.^36^ Several studies have suggested that the IOP-induced LC strains near the central retinal vessels (e.g., within the shadow regions) are smaller than further away.^12,28,29,31^ These are based on a similar rationale of the pre-mortem vasculature providing structural support to the adjacent tissues. Thus, although our method reduces the region available for analysis, it is necessary to avoid a potentially crucial bias.

Third, a reader may wonder if pre-mortem IOP variations could have affected the globe in a way that would affect the LC deformations measured post-mortem. We are confident that this is not the case. We selected the IOP levels to represent a wide range from low, but physiologic, to high, but still realistic. Nevertheless, these IOP levels are well within the range of IOPs that happens during eye rubbing^65^ and well below those expected to cause acute damage to the eye.^66^ Further, only for one of the animals (M2) we analyzed IOP variations in the same eye. In the other two animals (M1 and M3), we made our measurements in different eyes pre-mortem and post-mortem. This may help explain the larger difference in the IOP-induced LC strains between pre-mortem and post-mortem conditions for M1 and M3 than that for M2. However, this should not affect our conclusions, as previous studies in monkeys using histomorphometry have demonstrated that the contralateral eyes had much more similar deformations than unrelated eyes.^11^ While the assumption that the contralateral eyes have similar responses to IOP may apply to monkey eyes, it is still unclear how well it holds for human eyes. Some studies of enucleated eyes have reported good contralateral similarity in acute response to IOP,^13^ whereas others found a somewhat low contralateral similarity.^12^ Overall, we acknowledge that our comparisons for M1 and M3 rest on the assumption that contralateral eyes of an animal have similar acute responses to IOP.

Fourth, we did not manipulate ICP in the post-mortem condition. As a result, the translaminar pressure difference (IOP – ICP) post-mortem may be different than that pre-mortem. In addition, the extraorbital pressure may change, and the support from the extraocular muscles may be different post-mortem. Therefore, the changes in strains in the LC may be a combination of lack of blood pressure, change in ICP and extraorbital pressure, and altered muscle load. How exactly these changes contribute to the decreased LC strains post-mortem remains unclear and should be clarified in future studies.

Fifth, although we waited after pressure changes and between conditions to allow the system and tissues to stabilize (Figure 1), the interval necessary to fully converge remains unknown. Similarly, it is possible that in eyes tested more than once (M2) the first test affected the second one. There were constraints on the choice of interval length. On one hand, longer intervals after pressure changes and between conditions would help reduce potential effects of the transitions.

On the other hand, longer experiments may adversely affect image quality. The intervals chosen were selected based on the literature and our own preliminary work to balance these constraints.

In the study, we compared the IOP-induced LC deformations pre-mortem and post-mortem in five eyes of three monkeys, and the comparison results were consistent through the eyes we studied. It is worth reminding the reader that the experiment and analysis were not designed to determine whether our observations in these monkey eyes will extend to other eyes or monkeys. This cannot be determined from the small set of eyes we studied. The goal of our analysis was to obtain a robust set of findings for this particular set of eyes. Additional studies with more eyes and animals are necessary to apply our findings to the general population.

In summary, we have provided novel measurements of the LC deformations resulting from acute IOP increases immediately after death by exsanguination. We also measured the effects of IOP, allowing for a direct pre-mortem vs. post-mortem comparison. Overall, the acute IOP-induced LC deformations were smaller post-mortem than pre-mortem, but the trends in the nonlinear relationship with IOP were similar. Locally, deformations pre-mortem and post-mortem can be substantially different, with some regions being much smaller and some larger post-mortem than pre-mortem. Our results are expected to help improve the fundamental understanding of how LC biomechanics pre-mortem and post-mortem relate to one another, and how they depend on blood pressure.

## Acknowledgments

We thank Huong Tran for help throughout the project.

